# Cell adhesion marker expression dynamics during fusion of the optic fissure

**DOI:** 10.1101/2023.08.01.551419

**Authors:** Holly Hardy, Joe Rainger

## Abstract

Tissue fusion is a critical process that is repeated in multiple contexts during embryonic development and shares common attributes to processes such as wound healing and metastases. Ocular coloboma is a developmental eye disorder that presents as a physical gap in the ventral eye, and is a major cause of childhood blindness. Coloboma results from fusion failure between opposing ventral retinal epithelia, but there are major knowledge gaps in our understanding of this process at the molecular and cell behavioural levels. Here we catalogue the expression of cell adhesion proteins: N-cadherin, E-cadherin, ZO-1, and the EMT transcriptional activator and cadherin regulator SNAI2. We find that fusion pioneer cells at the edges of the fusing optic fissure have unique and dynamic expression profiles for N-cad, E-cad and ZO-1, and that these are temporally preceded by expression of SNAI2. This highlights the unique properties of these cells and indicates that regulation of cell adhesion factors is a critical process in optic fissure closure.

## INTRODUCTION

Tissue fusion underpins normal vertebrate embryonic development for a range of organs and tissues, and its dysregulation results in multiple structural birth defects, such as spina bifida, cleft palate, heart defects, and hypospadias [1–3]. Fusion events also display molecular and cell-behavioural overlap with wound-healing and tumour metastases, including the activation of EMT like processes [1,4], of which a critical initiating step is the decoupling of cell-cell adhesion complexes [5].

Ocular coloboma (OC) is a structural eye malformation defined by the failure of optic fissure closure (OFC) – an epithelial fusion event in the developing ventral retina [6–8]. OC is clinically characterised by a persistent gap at any region along the proximal distal axis of the ventral retina, and can affect the iris, retina, or optic nerve, and results in a range of impacts on vision [8]. Genetic diagnosis rates are low: up to 80% of coloboma patients currently do not have an identified causative mutation[9]. Furthermore, few loci have been identified with recurrent mutations across unrelated patients [10,11]. We previously illustrated that the developing chicken eye provides an accurate model for understanding the cell behaviours and molecular regulation of OFC [12,13]. Fusion begins at Hamburger Hamilton [14] embryonic chicken stage HH28 (∼ incubation day 6; **Figure 1**), when the two optic fissure margins (OFMs) come into close contact. By HH30, and over the following period to HH34 (∼day 8.5), chicken fissures fuse bi-directionally along the proximal-distal axis and display the defining features features of active epithelial sheet fusion [1]: open regions, leading edge apposition, cell mixing at a fusion plate, and a fully fused seam (**Figure 1**) [12]. During this process, a discrete population of cells at the optic fissure margins actively mediate fusion (OFM) [11– 13,15,16]. These *Pioneer cells* [12] are positioned between the developing retinal pigmented epithelium and neural retina (**Figure 1**), and in chick, zebrafish, mice, and humans, these undergo a partial EMT as they initiate the fusion process [12,15–17]. In human OFC, pioneer cells lose their typical epithelial morphology, express vimentin, and de-laminate from the neuroepithelium [17]. In chick, this is associated with subsequent apoptosis [12], and in zebrafish there is a highly-transient rearrangement of N-cadherin observed at the fusion plate [15]. In addition, EMT and markers of cell-adhesion have been implicated in OFC through ontology enrichment and OFC transcription profiling studies [12,17], however no definitive markers for EMT or cell adhesion have been qualitatively assessed *in situ* during the active fusion process.

**Figure 1.**
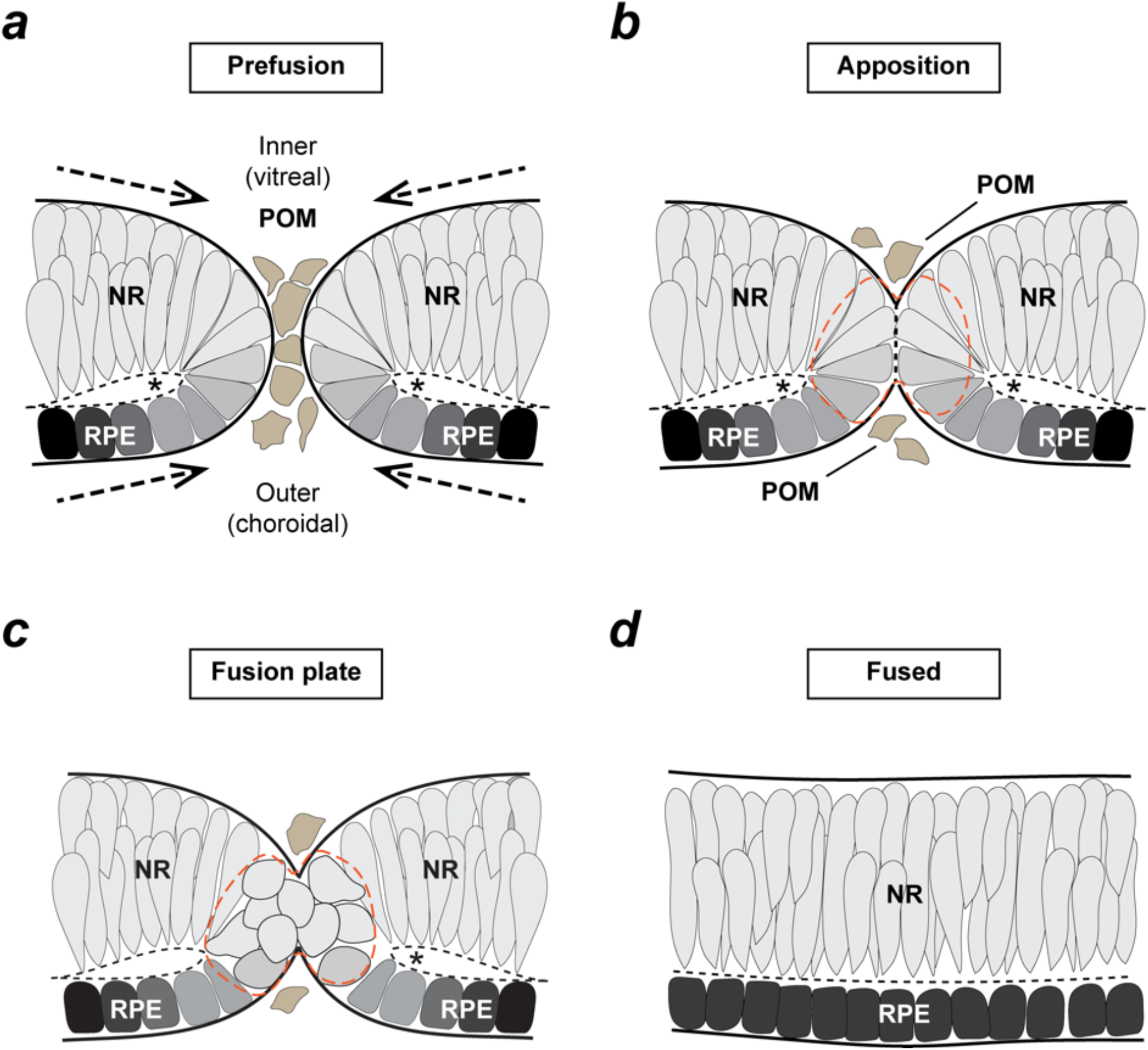
Schematic of fusion progress during vertebrate optic fissure closure. (***a***) In the ventral vertebrate retina at pre-fusion stages the optic fissure margins (OFM) approach each other as the optic cup grows (arrows). The OFM environment is composed of epithelial cells from the retinal pigmented epithelium (RPE) and neural retina (NR) and is interspersed with cells from the periocular mesenchyme (POM). Asterisks depict the folding point. The inner (vitreal) region is of the OFM is in the dorsal aspect of the fissure, and the outer region (choroidal) is ventrally positioned. (***b***) Fissure margins are in apposition immediately prior to fusion. Basement membranes form both edges of the fissure are in direct contact, and POM cells are displaced. (***c***) In the fusion plate, epithelial cells at the interface of retinal pigmented epithelium (RPE) and neural retina (NR) decouple from their neighbouring epithelial cells and mix to mediate active fusion - these are the fusion *pioneer cells* (red hatching). (***d***) In the nascently fused OFM, there are now fully organised and distinct RPE and NR layers surrounded by newly configured basement membranes at their inner and ventral layers. Figure adapted from Chen, Moosajee, and Rainger [20].

Here, we describe the expression of the key type I classical cadherin proteins N-cad and E-cad, and the tightjunction protein ZO-1 during the progression of fusion in the chicken embryonic retina epithelia. We find that pioneer cells have unique expression profiles for these cell adhesion molecules compared to the surrounding epithelia, and that this is dynamic during the fusion process. We also found pioneer-cell specific expression of the EMT and cadherin regulator SNAI2 during fusion immediately before these changes. Our data suggest these factors may combine to alter localised cell-adhesion properties and drive the cell behaviours required for epithelial fusion during OFC.

## RESULTS

### The tight junction marker ZO-1 is reduced in pioneer cells

We found the tight-junction cell-adhesion junction marker ZO-1 (encoded by the *TJP1* gene) was absent from cells at the pioneer cell region in pre-fusion OFM (**Figure 2a**). These OFM edges were still enveloped by basement membrane, as shown by laminin distribution (**Figure 2a**). In contrast, we observed ZO-1 positive staining in the adjacent non-fusing epithelial cell regions of the neural retina and RPE, with the strongest signal observed at the apical regions of these cells. This ZO-1 distribution is consistent with the presence of tight junctions (TJs) at the apical regions of NR and RPE. At the fusion plate (**Figure 2b**), where the pioneer cells were mixing across the connected OFM (as seen by displacement of laminin), cells were still negative for ZO-1 immunostaining. However, in the fused region of the OFM seam (**Figure 2c**), at a distance ∼100 um from the fusion plate ZO-1 staining was restored in the midline with ZO-1 distribution analogous to the adjacent neural retina and RPE regions that did not take part in fusion, indicating the establishment of newly-formed TJs in this region. This data shows that pioneer cells have reduced ZO-1 as they mediate the fusion process, suggesting the TJs in this region are disassembled as part of a process of cells decoupling from their neighbours to enable mixing and migration at the fusion plate.

**Figure 2.**
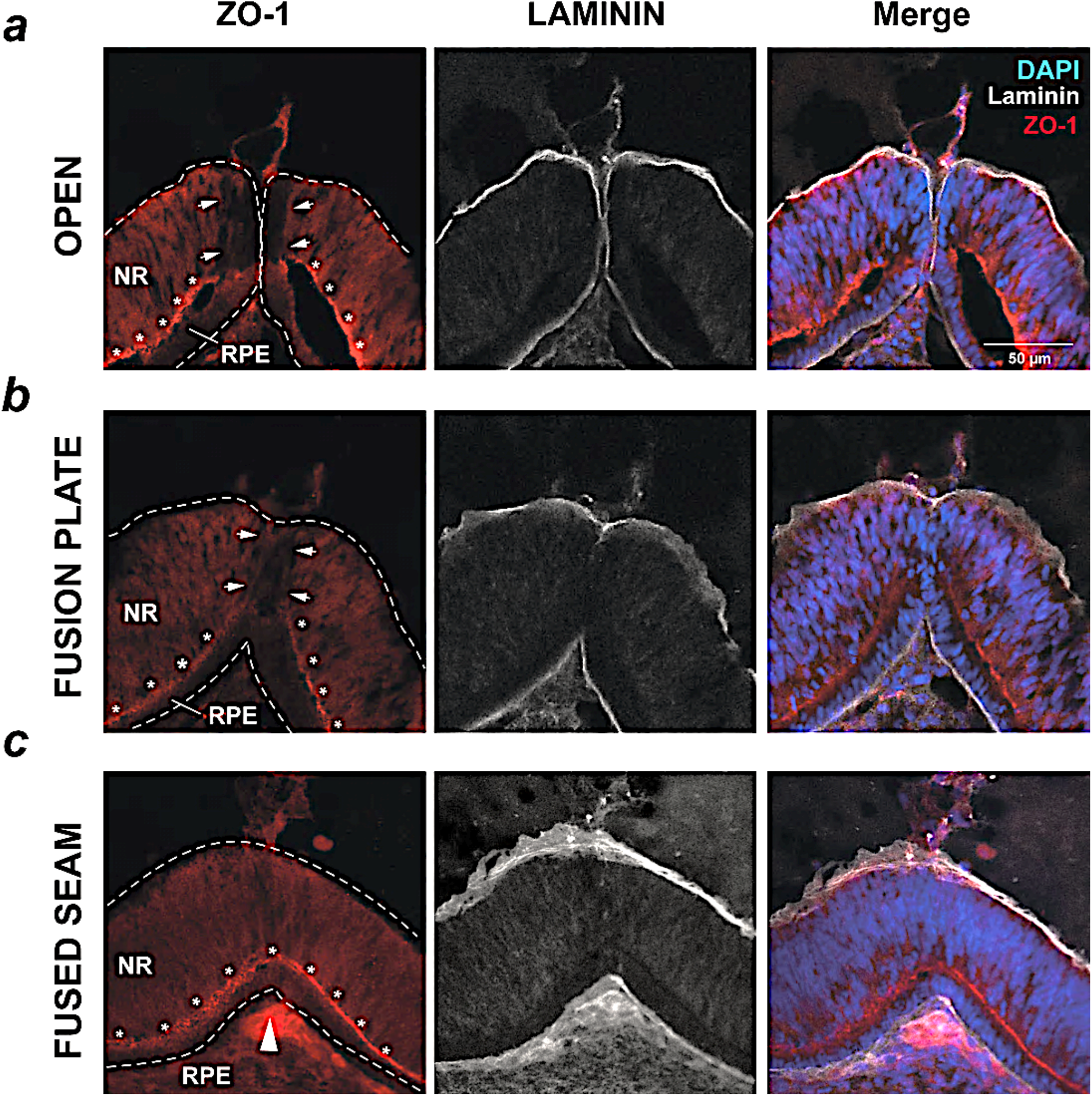
ZO-1 displays distinct expression during fusion. (***a***) Immunofluorescence staining for the cell junction marker ZO-1 and Laminin during active fusion (HH30) in the OFM revealed absence of ZO-1 expression in the distal Bp of the open fissure in the pioneer cell region (arrowheads). Strong ZO-1 signal was observed in the region where NR and RPE meet at their apical edges (astersiks). (***b***) ZO-1 expression was still reduced in the nascently-fused fusion plate where cells from the opposing margins had mixed. (***c***) Positive staining for ZO-1 was observed in the fusion seam (large arrow depicts midline) at ∼100 μm from the fusion plate and in the now continuous region where the RPE and NR cells are in contact at their apical edges (asterisks). Scale bar = 50 μm.

### Pioneer cells display heterotypic cadherin expression

To investigate the initial process of the EMT-like phenotype observed in pioneer cells, we performed immunostaining for the type I family of cadherins E-cad (CDH1) and N-cad (CDH2) at the optic fissure in HH30 chick eyes, when the OFM has all stages of active fusion available for analysis. We found that N-cad was strongly localised to the neural retina (**Figure 3a**) and absent from RPE and periocular mesenchyme. In the pioneer cell region, we observed some positivity for N-Cad, but at a lower intensity than in the adjacent NR. This reduced N-cad was present throughout the pioneer cell region during active fusion, and persisted in the fusion plate of the OFM where pioneer cells were actively mixing across the now fused epithelia (**Figure 3b**). However, in the midline of the nascently fused seam region of the same retinas, we observed that the N-cad signal was indistinguishable from the adjacent regions of epithelia (**Figure 3c**).

**Figure 3.**
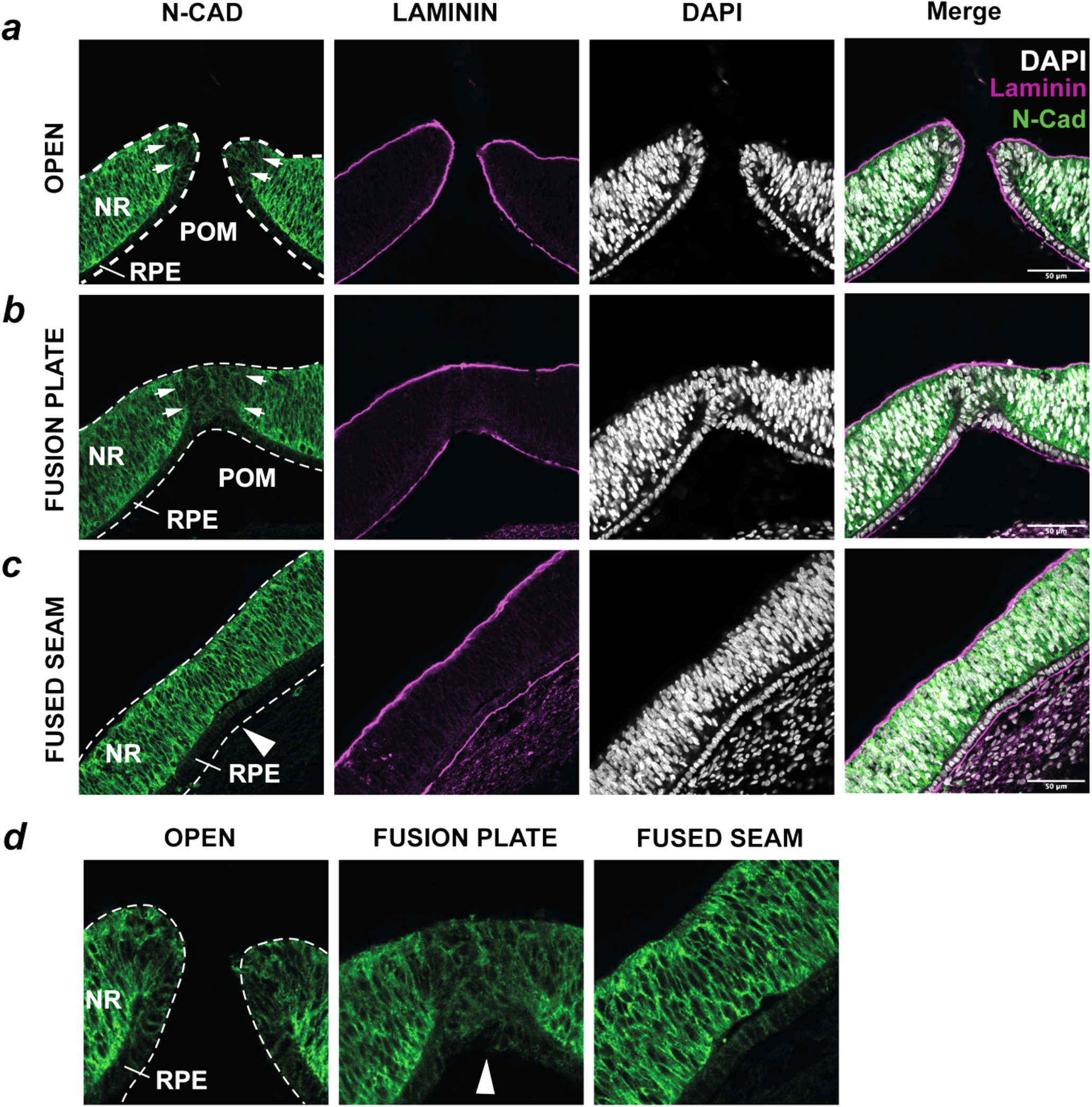
N-cadherin displays distinct expression during fusion. (***a***) Immunofluorescence staining for N-Cad/CDH2 and Laminin during active fusion (HH30) OFM revealed a localised reduction of N-Cad at the distal Bp regions of the open fissure (arrowheads). (***b***) Reduced N-Cad was persistent in the nascently-fused fusion plate. (***c***) Positive staining for N-Cad was observed in the fusion seam at ∼100 μm from the fusion plate (Large arrow depicts midline). (***d***) Enlarged views of *a-c* illustrating N-Cad expression at the midline during progression of fusion. Scale bar = 50 μm.

We then asked if we could see a pioneer cell specific expression pattern with E-Cad in stage-matched OFMs (HH30). We found that the E-cad expression pattern was reciprocal to that of N-cad, with strong expression observed in the RPE but no expression detected in the NR (**Figure 4**). Surprisingly, we also detected some E-cad expression extending from the RPE into in the pioneer cell region of the OFM. Similar to N-cad, this pioneer cell expression was at a lower intensity than the signal in adjacent cells, but the expression persisted throughout the active fusion stages (**Figure 4a,b**) but then was markedly reduced in the fused seam (**Figure 4c,d**). We did not detect E-cad expression in the periocular mesenchyme (POM). Using existing transcriptomic data from published datasets, we asked what other cadherins may be expressed in the OFM. In bulk RNAseq performed on dissected chick OFMs, *CHD4*, the gene encoding R-cadherin (R-cad), displayed fissure specific enriched expression [12]. Our qRT-PCR from freshly detected optic fissure samples at HH28 confirmed this enrichment for *CDH4* expression compared to dorsal retina tissue (**Supplemental Figure 1**). We were unable to successfully detect R-cad protein expression in chick samples with commercially available antibodies but were able to detect *CDH4* mRNA using fluorescence in situ hybridisation (**Supplemental Figure 1**). *CDH4* was specifically expressed in pioneer cells in pre-fusion (HH28) and at active fusion stages (HH30) (**Supplemental Figure 1**). Thus, these experiments indicate that pioneer cells have their own unique heterotypic cadherin profile compared to the adjacent epithelia, which constitutes overlapping low levels of both N-cad and E-cad at the protein level, and *CDH4* expression at the mRNA level.

**Figure 4.**
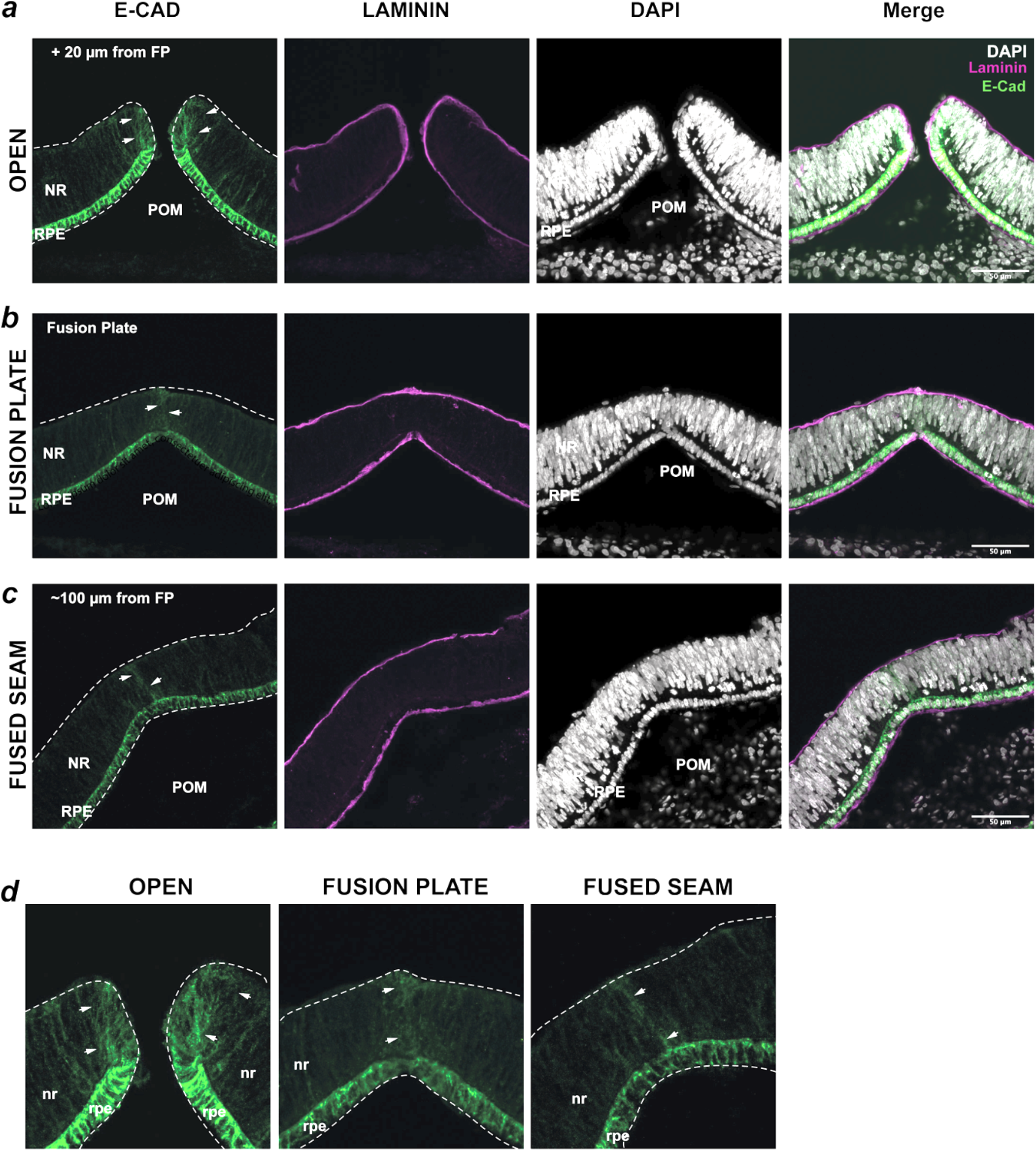
E-cadherin displays distinct expression during fusion. (***a***) Immunofluorescence staining for E-Cad/CDH1 and Laminin during active fusion (HH30) in the OFM revealed strong expression in the RPE that extended its domain into the distal Bp of the open fissure in the pioneer cell region (arrowheads). (***b***) Reduced E-Cad was observed in the nascently-fused fusion plate. (***c***) Residual staining for E-Cad was observed in the fusion seam at ∼100 μm from the fusion plate. (***d***) Enlarged views of *a-c* illustrating E-Cad expression at the midline during progression of fusion. Scale bar = 50 μm.

### SNAI2 is specifically expressed in pioneer cells during early OFC

Finally we sought to identify what factors may influence this heterotypic cadherin expression in the OFM. Our qRT-PCR analysis of the EMT transcriptional activators *SNAI1* and *SNAI2* showed increased expression of *SNAI2* in chicken pre-fusion OFMs compared to dorsal retina, whereas *SNAI1* expression levels showed no difference between these tissues (**Supplemental Figure 1**). We then performed both fluorescence *in situ* hybridisation and immunofluorescence analyses for SNAI2 and revealed expression at both the mRNA and protein levels in the pioneer cell region of the OFM prior to fusion (**Figure 5a; Supplemental Figure 1**). This expression was transient, as we were unable to detect SNAI2 expression in day 4 (HH22) OFM edge cells (approximately 48h before fusion) or in pioneer cells at the fusing and fused OFM (HH30) (**Figure 5b** and **Supplemental Figure 1**). SNAI2 expression at the protein and mRNA levels was detected in the POM surrounding the OFM and dorsal retina throughout all stages analysed, consistent with the migration of neural crest cells in these locations. Thus, SNAI2 expression in pioneer cells may provide a molecular link to the regulation of cadherin levels in these cells leading up to fusion.

**Figure 5.**
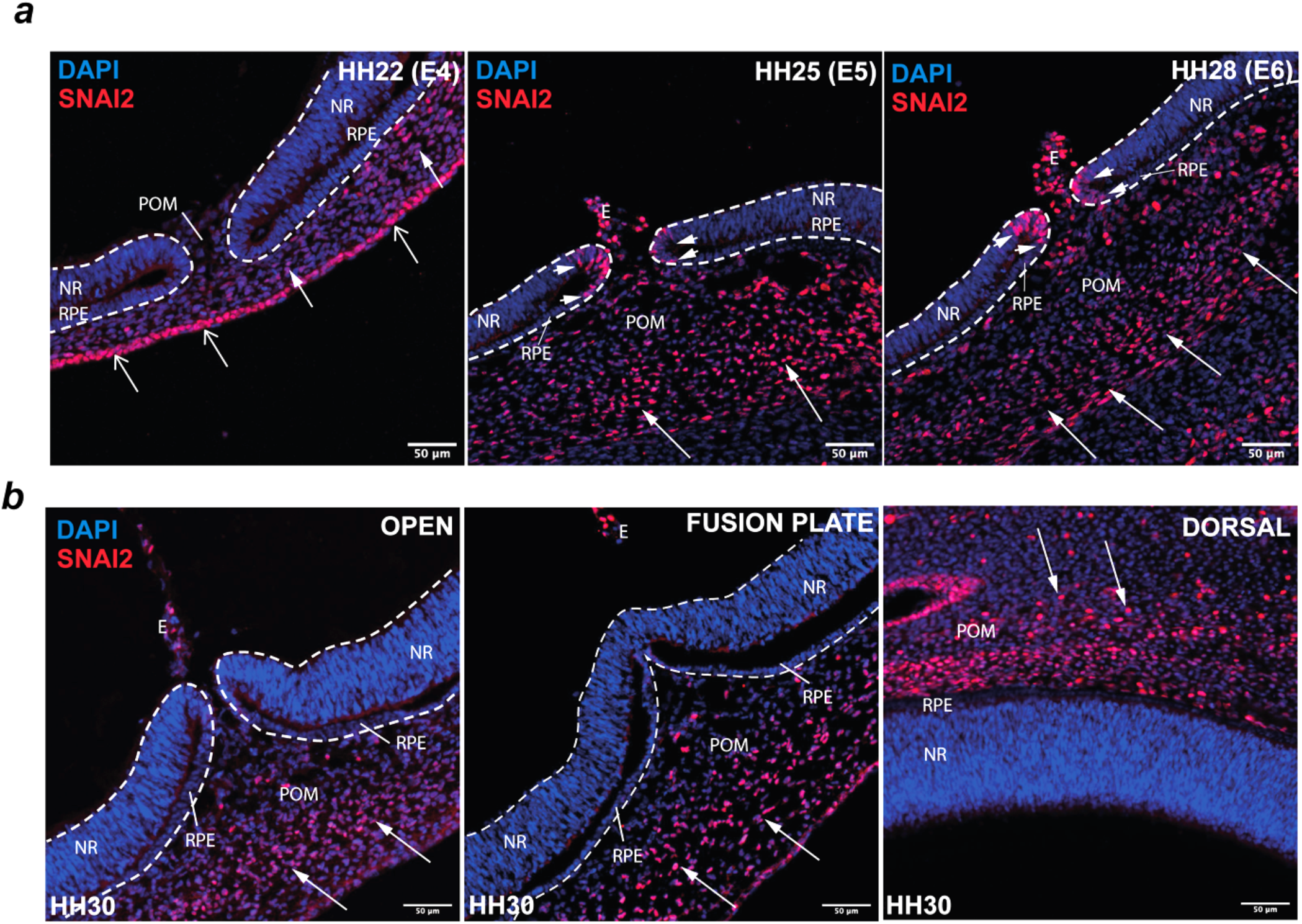
SNAI2 expression was observed in pioneer cells prior to fusion. (***a***) Immunofluorescence analyses showed SNAI2 was localised to the pioneer cell domains (arrowheads) in the pre-fusion stage OFM at HH25 and HH28 but was absent from the fissure margins in the earlier optic cup at HH22 (E4). Strong expression was observed in all stages in the POM (filled arrows) and was observed in the surface ectoderm (open arrows) at HH22. SNAI2 was also detected in the ectoderm of the developing pecten (E). (***b***) No SNAI2 was observed in the pioneer cell domains at HH30 in the open OFM and in the fusion plate. SNAI2 was consistently detected throughout the POM (filled arrows), including in dorsal eye tissue. Scale bars = 50 µm.

## DISCUSSION

Determining the precise molecular signatures of the cells that directly mediate tissue fusion during optic fissure closure has been challenging due to the small size of this population and their transient temporal existence. Here, using immunofluorescence and fluorescence *in situ* hybridisation profiling we provide qualitative evidence that these are temporally distinct cell populations, and show they have dynamic and specific expression for the cell-cell adhesion molecules E-cad, N-cad, and ZO-1. We also show that the expression dynamics of these cell-cell adhesion factors are temporally preceded by the transient expression of SNAI2. These novel findings will inform future studies to understand some of the mechanisms that mediate cell behaviours during fusion in the optic fissure.

Cell-cell cohesion is typically mediated by homophilic adhesive cadherin binding, with cells tightly joined to neighbouring cells expressing the same type I cadherin via selective adhesion between cell types[18]. Consistent with this function, we observed strong N-cad expression in the neural retina, and reciprocally, strong E-cad expression in the retinal pigmented epithelia. These two retina epithelial tissues are relatively stable, and during the remaining phases of eye development will either undergo further differentiation into photoreceptor cell types (neural retina) or will mature as a single epithelial layer (RPE) [7,19]. Thus, stable cell-cell adhesion supports their further development. In contrast, epithelial cells at the edges of the OFM must decouple from their immediate neighbours to actively facilitate fusion [20], a process that presumably would require a loss of cell-cell adhesion. Here, we provided supportive evidence for this through observed reduction of ZO-1 expression in pioneer cells at pre-fusion stages and during active fusion.

Our data revealing the heterogeneous cadherin profile of these cells suggests that cadherin dynamics may promote epithelial decoupling during OFC, but this expression profile could also have a role in defining pioneer cell boundaries during retinal development. The observation of overlapping E-cad and N-cad protein expression at pioneer cell surfaces is in line with the “differential adhesion hypothesis” (DAH) that proposes that distinct cell types can be sorted because of tissue surface and interfacial tensions as a function of their cadherin expression [21,22]. In this context, RPE and NR are defined by their expression of E-cad and N-cad, respectively, whereas pioneer cells are defined by their heterogeneous N-cad, E-cad and Rcad cadherin expression. In addition to forming a buffer between NR and RPE, this heterotypic cadherin expression may be sufficient to determine pioneer cell identity, as direct crosstalk among classical cadherins has recently been shown to form heterotypic connections [23,24].

Our favoured hypothesis is that the heterotypic cadherin presentation acts as a driving mechanism for reduced cell-adhesion in these cell populations, enabling cell decoupling, mixing and intercalation across the margins to initiate fusion. This process may also trigger localised changes and breakdown of the ECM and overlying basement membranes, but further work is required to provide direct evidence for this.

Intriguingly, both RNAseq [12,17] and our data here (**Supplementary Figure 1**) suggest that additional type I cadherins are likely to be expressed in pioneer cells, such as *CDH6* and *CDH4*/Rcad, and this may further increase the diversity of heterotypic cadherin presentation at cell-surfaces and affect cell-cell adhesion dynamics. Whether these different type I cadherins directly and functionally interact within the pioneer cell population remains unknown, but there is existing evidence that this may be the case [23,24], including a recent study that found both co-expression and molecular interactions between heterotypic E-cad and N-cad in the developing chicken neural tube [24]. A further possibility is that the *level* of cadherins expressed is important in regulating pioneer cell adhesion dynamics, as the number of cadherin molecules presented on cell surfaces is directly related to cell-cell adhesion properties [21]. Thus, to fully understand how pioneer cells are de-epithelialized during the early stages of fusion, future work in this area should determine the full complement and levels of adhesion molecules expressed in these cells, and their functional interactions, including the downstream effects of heterotypic cadherin co-expression. This current study therefore provides a framework for ongoing analyses into cadherin roles during OFC by identifying the presence of heterotypic cadherin expression in pioneer cells.

How cadherin expression is regulated in this small and transient cell population is currently unknown. The transcriptional repressor SNAI2 is a mediator of EMT (reviewed in [25]) and can both directly and indirectly regulate the expression of cadherins, including E-cadherin via interaction with the E-cadherin promoter [26,27]. SNAI2 was therefore a strong candidate for this function in pioneer cells at the OFM and so we assessed SNAI2 expression at both the mRNA and protein levels. We found that SNAI2 was specifically expressed in pioneer cells, but only at pre-fusion stages and not during active fusion or in the fusion plate. As the heterotypic expression of cadherins in pioneer cells was only observed at pre-fusion and active fusion stages, but was resolved to cell type (NR or RPE) in the nascently-fused retina, this suggests that if SNAI2 does directly regulate the precise balance of cadherin expression in pioneer cells, it is likely to function in this role principally to set the pioneer cells up to drive fusion, rather than acting as a continually active factor in this process. The functional requirement for SNAI2 during OFC, and confirming its role in balancing cadherin expression, will be an important next step in understanding pioneer cell behaviours and their regulation, warrants further work.

## METHODS

### Chicken embryo processing

Wild type Hy-Line chicken eggs were collected and incubated at 37°C and collected for analysis of OFC fusion between 5 – 7 days. Active fusion begins at HH28/Day 6 and is fully complete by HH32/Day 8 [12]. All embryos were staged according to Hamburger and Hamilton criteria and for fusion progression based on Hardy et al. For each assay, a minimum of 3 separate eyes from different embryos were used as replicates. We did not sex embryos, but found no variation in our observations between sample replicates. Embryos were rinsed in PBS (pH 7.0) on ice for 5 minutes then fixed in 4 % paraformaldehyde for 4 hours at 4°C with gentle agitation. Embryos were then rinsed three times in PBS (pH 7.0) for 15 minutes on ice and then incubated in 10 % sucrose PBS (pH 7.0) overnight at 4°C with gentle agitation. Samples were embedded in Neg50 embedding media (Epredia) and snap frozen as either whole embryos (HH22-HH28/day 4-6) or, for HH30 (E7) samples, ventral retinas were dissected only, then stored at 80°C until further use. Sections were cut on a Leica CM1900 cryostat at 14 µm and sections were attached onto Superfrost Plus glass slides (Epredia) air dried for 1 hour at room temperature, then stored at 80°C until further use.

### Immunofluorescence

Slides were rinsed in PBS for 5 min at room temperature, then incubated in Block #1 (2 % bovine serum albumin, 0.25 % Triton-X-100 in PBS, pH 7.0) for up to 5 hours at room temperature. Primary antibodies were diluted as stated below in Block #2 (0.1 % bovine serum albumin, 0.05 % Triton-X-100 in PBS, pH 7.0) applied to slides and then incubated overnight at 4°C. Slides were then rinsed three times in PBS for 5 min at room temperature and then PBS was replaced with secondary antibodies diluted 1:1000 in Block #2 and incubated for 1.5 hours at room temperature. Slides were then rinsed three times in PBS for 5 min at room temperature and PBS was removed and slides were mounted in Fluorsave (Millipore; #345789) and allowed to cure overnight in a dark chamber. Primary antibodies used and their working dilutions were rabbit anti SNAI2 (C19G7) (Cell Signalling Technology #9585) at 1:100; mouse monoclonal anti E-Cadherin antibody (BD Biosciences #610181) at 1:2500; anti ZO-1 rabbit polyclonal antibody (Thermo #40-2200) at 1:100 dilution, anti Laminin (DSHB #3H11) at 1:20, and mouse monoclonal anti-N-Cadherin antibody (Thermo #C3865) at 1:100. We used Alexa Fluor IgG (H+L) Cross-Adsorbed fluorescent secondary antibodies (647 nm, 594 nm and 488 nm) (Thermo). For nuclei staining, we used DAPI (diamidino-2-phenylindole; Thermo #15105118) at 1 ug/mL incubated with secondary antibodies.

### Fluorescence in situ hybridisation

For *SNAI2* RNA fluorescence in situ hybridisation we used RNAScope (ACD, Biotechne) following the RNAscope Multiplex Fluorescent V2 Assay protocol on 14 µm cryosections prepared as above and processed as described previously [12,28] with the probe ID #1172211-C1. This probe was designed to target *Gallus gallus* snail family transcriptional repressor 2 (*SNAI2*) transcript variant X1 mRNA [NCBI GeneID:432368].

### Imaging

All imaging was captured on a Zeiss LSM 880 AxioObserver confocal microscope using Plan-Apochromat 20x/0.8 M27, Plan-Apochromat 63x/1.4 Oil DIC M27, or Plan-Apochromat 40x/1.3 Oil DIC UV-IR M27 objectives and Zen Black image software (Zeiss). Images were saved as .czi files and then opened in imageJ and converted to jpeg format.

### Institutional Animal Review Board Statement

Chickens were maintained according to institutional regulations for the care and use of laboratory animals under the UK Animals Scientific Procedures Act. All chicken embryology work was carried out on embryos prior to 2/3 gestation, and no regulated procedures were performed.

### Open access

For the purpose of open access, the author has applied a CC-BY public copyright licence to any Author Accepted Manuscript version arising from this submission.

## Contributions

Conceptualization - JR; Data curation – JR & HH; Formal analysis - JR & HH; Funding acquisition - JR; Investigation - HH & JR; Methodology – HH & JR; Project administration - JR; Roles/Writing - original draft - JR; and Writing - review & editing – HH & JR.

## Acknowledgments

We would like to thank Roslin Institute staff at the Greenwood Building for chicken husbandry and the Bioimaging and Flow Cytometry Facility for help with microscopy.

## List of Abbreviations

OFC: Optic fissure closure
OFM: Optic fissure margin
OC: Ocular coloboma
RPE: Retinal pigmented epithelium
NR: Neural retina
POM: Periocular mesenchyme

## SUPPLEMENTAL INFORMATION

**Figure S1.**
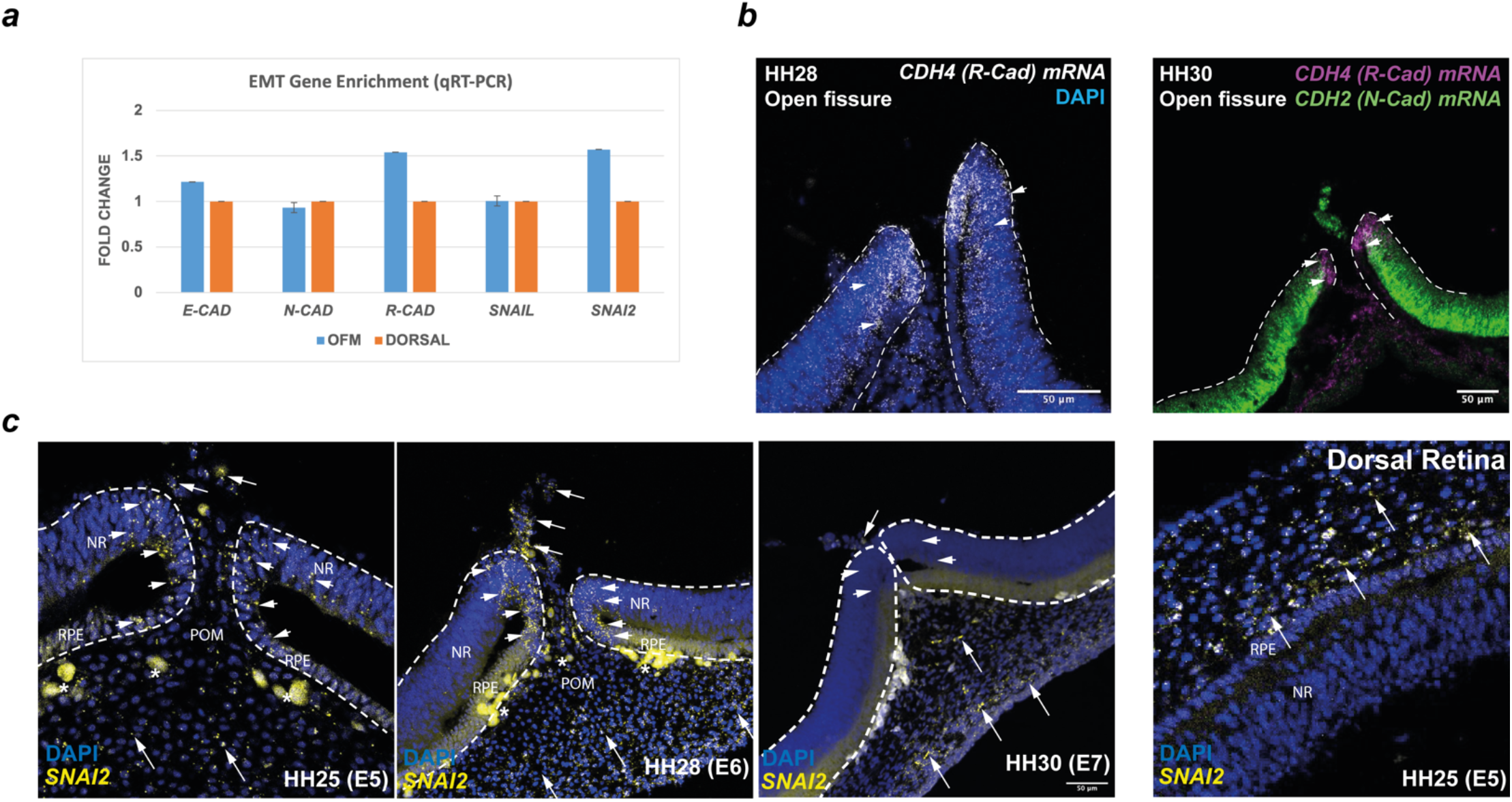
(***a***) qPCR RT analysis of *E-cad, N-cad, Rcad, SNAI1*, and *SNAI2*. (***b***) Fluorescence *In Situ* hybridisation using HCR shows enrichment for *Rcad* (*CDH4*) mRNA expression in the pioneer cell domain (arrowheads) of the fissure margin at HH28 and HH30. (***c***) Fluorescence *In Situ* hybrid isation using RNAscope with a probe for *SNAI2* mRNA shows expression in pioneer cells at the edges of the fissure margin (arrowheads) and in the POM (full arrows) in pre-fusion stages (HH25/E5 and HH28/E6). No NR/RPE SNAI2 expression was detected in dorsal retinal tissue but was observed in dorsal POM.

### Supplemental Methods

#### HCR Fluorescence in situ hybridisation

For N-cadherin (*CDH2*) and R-cadherin (*CDH4*) mRNA detection we used HCR RNA fluorescence in situ hybridisation according to the manufacturer’s instructions (https://www.molecularinstruments.com/). Briefly, embryos were developmentally staged, and whole optic fissures were dissected from the ventral eye and fixed in 4% PFA overnight. Fissure samples were processed (n>3 per run) for HCR then prepared as cryosections as described above. Gene specific probes were designed by Molecular Instruments: Chicken N-Cadherin (CDH2; NM_001001615.2) used with HCR Amplifier B1 [B1-488 nm]; Chicken R-Cadherin (CDH4; NM_001004391.2) used with HCR Amplifier B3 [B3 – 647 nm].

#### qRT-PCR

Optic fissure tissue and dorsal eye were dissected from wild type HyLine chicken embryos at HH28. Pools of either fissure or dorsal retina were collected (n = 10 per pool). 3x pools were prepared in total for each tissue type. Total RNA was extracted using Trizol reagent (Thermo) and assessed for quality by agarose gel electrophoresis. 1 µg total RNA was used in 1^st^ strand cDNA synthesis reactions using oligod(T) primers and Superscript III (Thermo) according to the manufacturer’s instructions. cDNA was used as template for qRT-PCR in 20 µl reactions using Luna Universal qPCR Master Mix from (New England BioScience). Oligonucleotide primers sequences are in **Supplemental Table S1**. Reactions were run on a Stratagene Mx3000p thermal cycler using a 2-step programme and critical threshold (CT) values were collected using MxPro software (Stratagene) and exported to Microsoft Excel to calculate fold enrichment relative to the *HPRT1* control by the comparative delta-delta-Ct method (2−ΔΔCt).

**Table S1.**
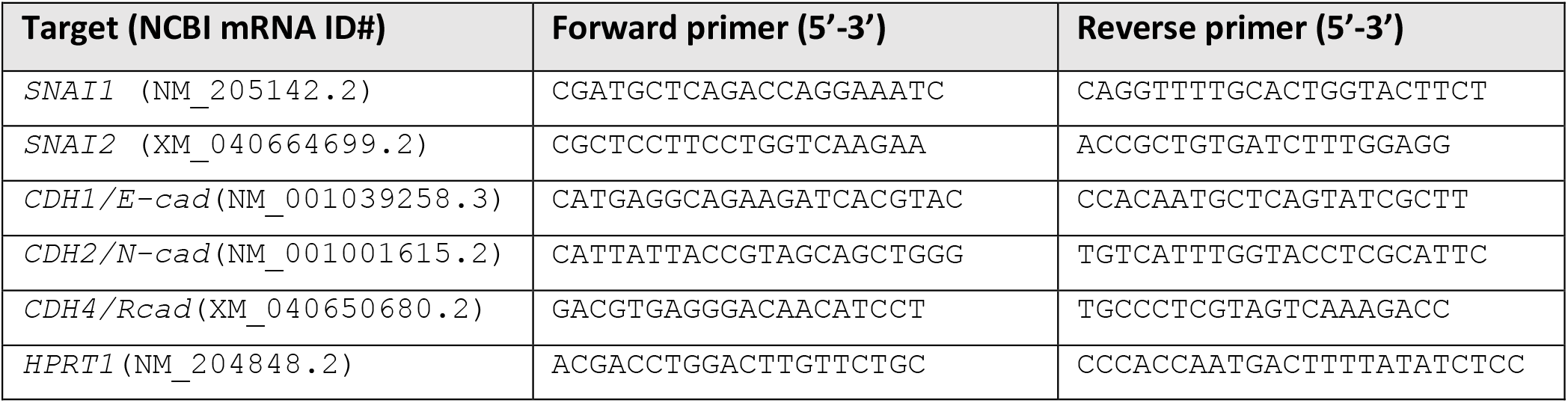
Primer sequences for qRT-PCR.

